# Structural bias in machine learning-guided peptide design

**DOI:** 10.64898/2026.05.06.721805

**Authors:** Victor Daniel Aldas-Bulos, Fabien Plisson

## Abstract

Machine learning continues to accelerate peptide and protein design through the rapid prediction and generation of sequences with desired characteristics. Many applications focus on predicting properties, functions, and structures, as well as generating point mutations and *de novo* designs. Nevertheless, many models prove less generalizable than initially claimed. Most predictors and generators are trained on sequential datasets, where imbalances can be addressed during preprocessing. In contrast, structural bias, a subtype of algorithmic bias arising from uneven representation of structural classes in training datasets, and the limitations of early protein structure predictors have frequently remained undetected and uncorrected. The recent surge in powerful protein structure prediction tools, such as the AlphaFold and RosettaFold series and their variants, now presents opportunities to mitigate this issue. We hypothesize that such structural sampling biases influence the downstream performance of ML models. Using antimicrobial peptides as a case study, we audited the structural biases in 16 state-of-the-art predictors for antimicrobial activity and tested whether structural information constrains their predictions. Our analysis revealed that models explicitly trained on sequential data still produce predictions biased by uneven fold representations and data leakage. These findings highlight the importance of integrating balanced structural data or implementing bias-mitigating strategies to develop agnostic models that maximize bioactive protein discovery and multi-objective optimization.

## 1. INTRODUCTION

Facial recognition systems that misidentify people with darker skin tones or women^1^, lending and credit scoring models that disadvantage minority or low-income communities^2^, and image generators that reproduce stereotypes embedded in the training data^3^ illustrate the pervasive nature of algorithmic bias in our society. These examples demonstrate that bias can emerge from human decision-making and the computational systems we design, often mirroring and exacerbating pre-existing inequalities. Algorithmic bias refers to systematic biases in the outputs of computational models or decision-making systems, arising from the data on which they are trained, the algorithms they use, or the contexts in which they operate.

The consequences of algorithmic bias in domains such as finance, law enforcement, and media production are increasingly recognized, prompting governments and private organizations to implement policies and guardrails that support the safe and responsible use of artificial intelligence (AI) technologies^4,5^. By comparison, the effects of algorithmic bias in the natural sciences, including computational biology, have received less attention. Over the past decade, the field of protein biology has benefited from machine learning (ML) methods for engineering, structure prediction, functional annotation, and *de novo* design^6–13^. However, because these models are trained on protein sequence or structural databases, they inherit the historical, methodological, and experimental biases embedded in these resources, shaped by decades of research funding priorities, experimental detection limits, taxonomic preferences, and domain-specific data availability^14^.

Recognizing and addressing such biases early is essential to developing robust, generalizable, and reproducible applications in protein engineering, drug discovery, and synthetic biology^15^. It may also present significant biosecurity risks. In line with these objectives, the Responsible AI × Biodesign initiative provides a global framework that outlines community values, guiding principles, and commitments to ensure sound AI-guided protein design practices^16^.

Early evidence of algorithmic bias in protein biology has emerged across sequence-, structure-, and function-prediction ML models. Sequence-based approaches rely on training datasets with phylogenetic and annotation biases that overrepresent certain model organisms, protein families, and conserved domains, thereby reducing model performance on underrepresented sequences^17–20^. Structure-based models exhibit performance variations due to imbalanced structure classes, fold-switching proteins, and intrinsically disordered regions in structural repositories^21–25^. Functional annotation models propagate historical misannotations from homology-based methods, overlooking novel functions^26–28^. These biases can distort predictions and generations, narrow the exploration of the sequence-function landscape to dominant protein classes, and limit the discovery of rare folds^29–31,24^.

To examine structural bias, a subtype of algorithmic bias, we recently applied a set of protein structure predictors, including AlphaFold2^9^, to map the structural landscape of medium-to-large-sized datasets using the GRAMPA repository of 5,980 antimicrobial peptides^32^. Our structural mapping revealed that most peptides (65.1%) adopted loose helices, whereas fewer formed random coils (17.8%) or β-stranded or mixed structures (17.1%) – mimicking folds observed in X-ray crystallography and aqueous NMR solutions. Because many state-of-the-art antimicrobial peptide predictors are trained on GRAMPA or similar repositories^1^, we evaluated 16 models to assess whether the latent structural composition influences their predictions and to explore how structural features shape predictive bias.

## 2. MATERIALS AND METHODS

### 2.1. Datasets

#### GRAMPA

We obtained the peptide sequences from the GRAMPA repository (Giant Repository of AMP Activities), a robust database established in 2018 containing sequences of 5,980 peptides ranging from 5 to 50 residues.^33^ The GRAMPA repository and detailed information are available at https://github.com/zswitten/Antimicrobial-Peptides.

#### Non-GRAMPA

We retrieved 3,385 peptide sequences ranging from 5 to 50 residues from the UniProt database^34^. These sequences had no reported antimicrobial activity and originated from taxa known to produce antimicrobial peptides: Arthropoda (n=1,867), Mammalia (n=501), Violaceae (n=114), Amphibia (n=322), Rubiaceae (n=49), Cucurbitaceae (n=60), and Mollusca (n=472). The search excluded entries labeled as partial, putative, or predicted, and those annotated with keywords *Antimicrobial* [KW-0929], *Antiviral*, *Anticancer*, *Antibiotic* [KW-0044], or *Fungicide* [KW-0295], while restricting results to reviewed records (*reviewed: yes*).

### 2.2. Preprocessing

#### Structural predictions

We used PEP2D (https://webs.iiitd.edu.in/raghava/pep2d)^35^ to predict the secondary structures of all peptides. Each peptide is represented as a vector of three-state values (H: helices, E: extended strand/β-sheets, and C: coils) expressed as percentages. The collection of peptide structure predictions forms the corresponding structural landscape, depicted in a ternary plot.

#### Structural classes

We defined seven structural regions (or classes) using the three states (H, E, C) and arbitrary borders at 80:0:20, 20:0:80, and 20:80:0.

#### Removal

We removed duplicated sequences, those containing non-canonical or unknown amino acids (represented as X), and sequences common to both the GRAMPA and non-GRAMPA datasets.

#### Redundancy

We used the CD-HIT^36^ web server (https://github.com/weizhongli/cdhit-web-server) to remove highly redundant sequences from both datasets, grouping sequences with a 70% identity threshold and retaining only the most representative sequence from each group.

### 2.3. Model selection, evaluation, and interpretation

#### AMP predictors

We selected 16 predictive models for antimicrobial activity: *AMP scanner v2*^37^, *amPEPpy*^38^, *AMPlify* variants^39^, *CAMPr3* variants^40^, *DBAASP*^41^, *IAMPE* variants^42^, *iAMPpred*^43^, *PepNet*^44^, and *Sense the Moment* ^45^ – see **Table S1**.

#### Evaluating the model performances

Each model was evaluated as a binary classifier, with antimicrobial peptides (AMPs) assigned to class 1 and non-antimicrobial peptides (non-AMPs) to class 0. Depending on the architecture, the model outputs were interpreted as either discrete class labels, class probabilities (*e.g.*, *p*(sequence X | AMP) = 0.67), or both. For models that produced only class probabilities, sequences with values greater than 0.50 were labeled as AMPs. For each structural class, we computed the confusion matrix, recording the number of True Positives (TP; correctly predicted active AMP sequences), True Negatives (TN; correctly predicted non-AMP sequences as inactive), False Positives (FP; non-AMP sequences incorrectly predicted as AMPs), and False Negatives (FN; AMP sequences incorrectly predicted as non-AMPs). From these, we derived standard performance metrics, including accuracy, precision, specificity, sensitivity, and Matthews Correlation coefficient (MCC).

1. Accuracy, the fraction of correct predictions obtained by the model, defined as:

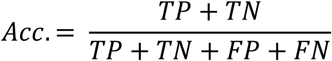
2. Precision, the fraction of positive predictions correctly identified:

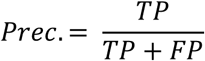
3. Specificity, the fraction of negative predictions correctly identified:

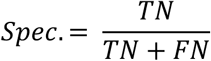
4. Sensitivity, the fraction of true positives correctly identified:

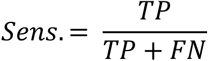
5. Matthews Correlation coefficient integrates information from all four elements of the confusion matrix, yielding a value between -1 and 1. Values of 1 and -1 indicate perfect and completely erroneous classification, respectively, whereas a value of 0 reflects random prediction.

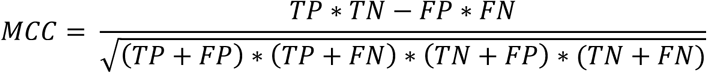

#### Embeddings and Dimensionality reduction of peptide datasets

We encoded the structures of the training sets *(amPEPpy*, *AMP scanner v2*, *AMPlify*, *PepNet*) and the external validation sets (GRAMPA and non-GRAMPA) using ProstT^46^, a sequence-structure encoder that generates 3Di tokens as proxies for three-dimensional representations. The resulting structural embeddings were projected onto a two-dimensional UMAP^47^ manifold optimized to separate the structural classes – see **Figure S6**.

## 3. RESULTS

Machine learning algorithms have become increasingly complex, gaining in performance but often at the expense of interpretability. Many existing models lack detailed descriptions of the training data and algorithms, which limits their reproducibility and contributes to their perception as “black boxes”. This limited transparency complicates the evaluation of potential biases and the robustness of model predictions. To address these concerns, explainable AI techniques have gained traction, and external (*post-hoc*) validation on independent datasets is a robust strategy for uncovering inherent biases.

In our study, we assembled two curated datasets to critically assess the performance and potential biases of existing binary classifiers for predicting antimicrobial peptide activity. The independent validation set comprised 5,980 antimicrobial peptide sequences from the GRAMPA repository^33^ and 3,385 non-antimicrobial peptide sequences from the UniProt database (see **Materials and Methods**). To explore the sequence-structure relationships with these sets, we predicted the secondary structure of all peptides using the PEP2D web server^35^. We then assigned each peptide to one of seven structural classes based on defined thresholds for the fractions of helix (H), extended strand/β-sheet (E), and coil (C).

### 3.1. Structural composition of the validation sets

The GRAMPA repository is enriched in α-helical and coiled structures, consistent with PEP2D and AF2 predictions, which were experimentally validated, as previously reported^32^. To assess structural differences from non-antimicrobial peptides, we predicted the secondary structures of the non-GRAMPA dataset. Initial analysis revealed a limited presence of β-sheet conformations in non-GRAMPA sequences. To better capture this structural class, we supplemented the dataset with 53 sequences explicitly annotated as β-sheet structures, bringing the total to 3,438.

We subsequently removed duplicate sequences and those containing non-canonical or ambiguous residues, yielding 2,511 representative non-GRAMPA peptides. Both GRAMPA and non-GRAMPA datasets were further subjected to redundancy reduction via CD-HIT^36^ clustering at 70% sequence identity. This process reduced GRAMPA to 1,731 sequences and non-GRAMPA to 1,185 sequences, thereby eliminating highly homologous peptides while preserving sequence diversity.

Importantly, this reduction did not substantially alter the distribution of structural classes across the datasets. The GRAMPA subset remained dominated by α-helices and coils, accounting for 64.6% of sequences, with 15.1% belonging to β-sheet/coiled structures and 14.1% to coils (**Table 1**). Conversely, the non-GRAMPA subset exhibited a more balanced structural composition, comprising 35.9% helices, 32.7% β-sheets/coils, and 25.6% coils. The structural landscapes of both datasets are illustrated in **Figure 1**.

**Figure 1.**
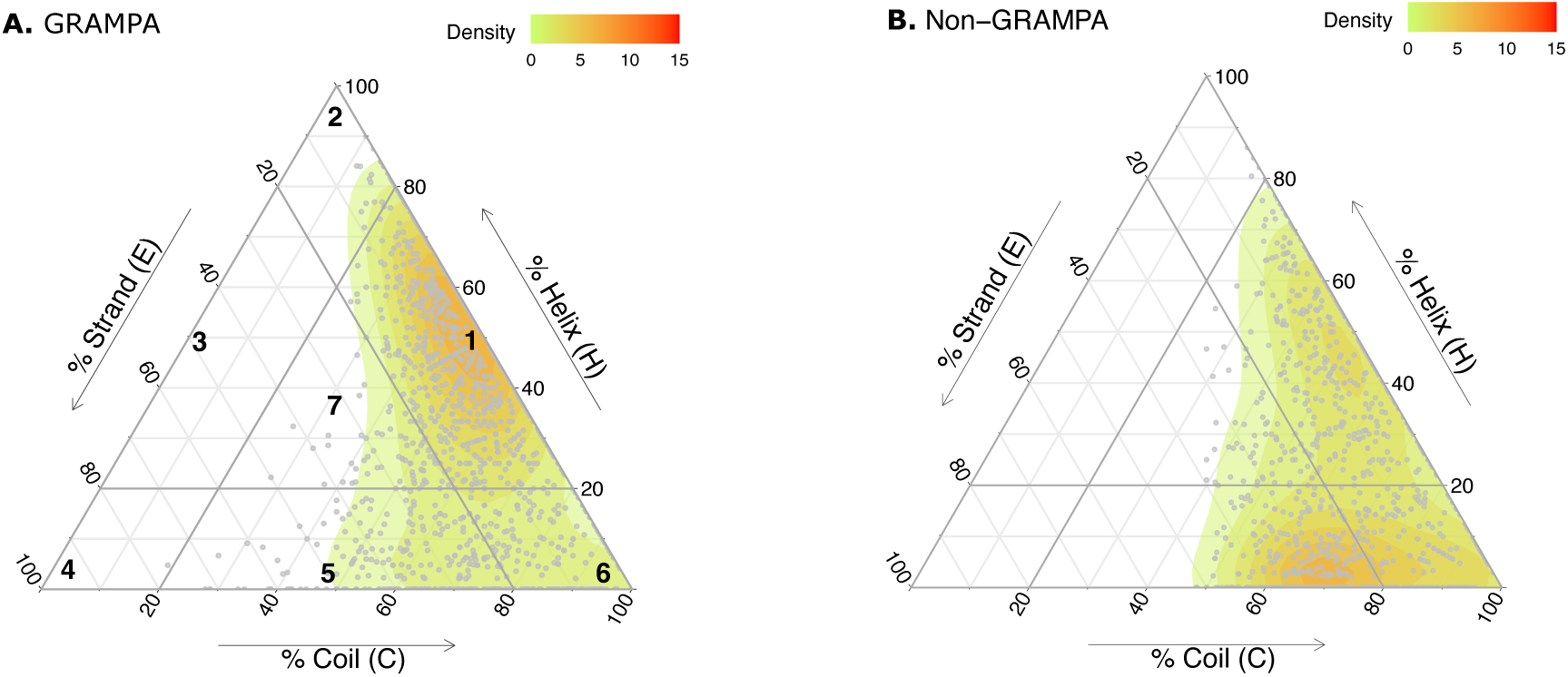
Predicted structural landscapes of (A) GRAMPA and (B) non-GRAMPA.

**Table 1.**
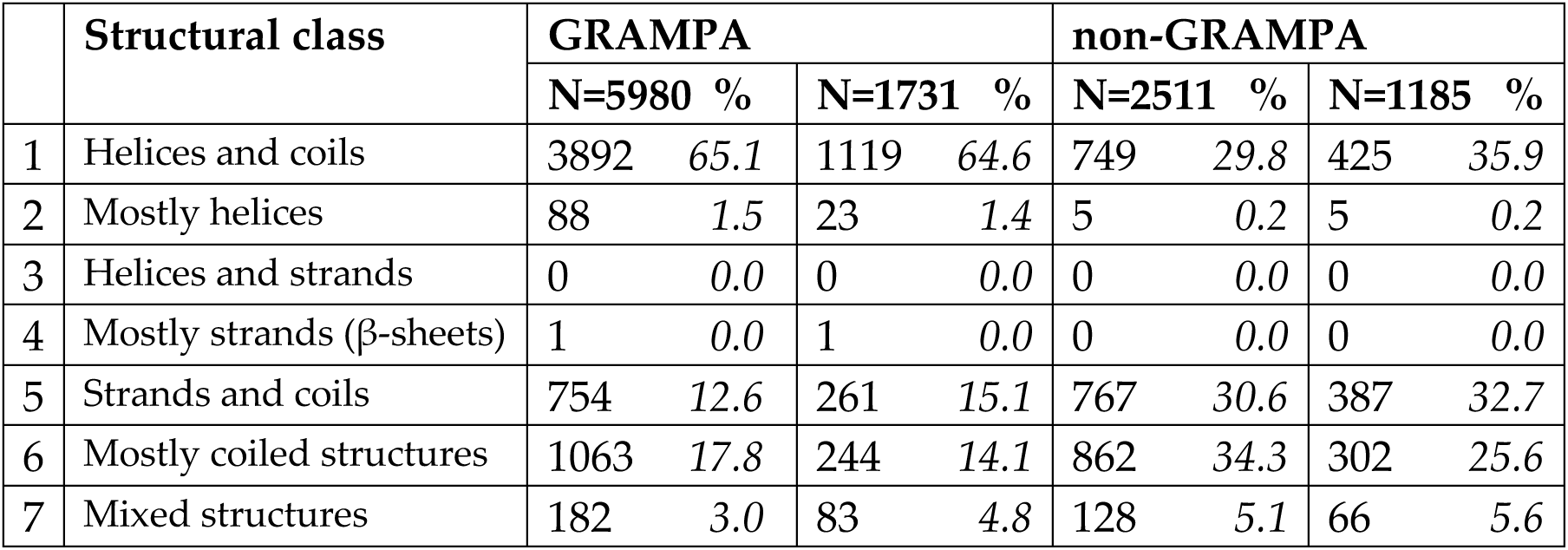
Distributions of predicted structural classes (1)-(7) for the GRAMPA and non-GRAMPA datasets using protein secondary structure predictor PEP2D.

### 3.2. Performance of AMP predictors across structural classes

To investigate whether the structural distributions of GRAMPA and non-GRAMPA subsets influence antimicrobial peptide (AMP) prediction models, we evaluated 16 state-of-the-art predictors published between 2016 and 2024 (**Table S1**). These models vary in machine learning architectures and training set sizes: *AMP scanner v2*^37^, *amPEPpy*^38^, *AMPlify* variants^39^, *CAMPr3* variants^40^, *DBAASP*^41^, *IAMPE* variants^42^, *iAMPpred*^43^, *PepNet*^44^, and *Sense the Moment* ^45^. Predictions were stratified into the four dominant structural classes, as shown in **Table 1**: helices and coils (1), strands and coils (5), mostly coils (6), and mixed structures (7). For each model and structural class, we calculated accuracy, precision, sensitivity, and specificity from the confusion matrices shown in **Figures 2A-D** and **Figures S1-S4.**

**Figure 2.**
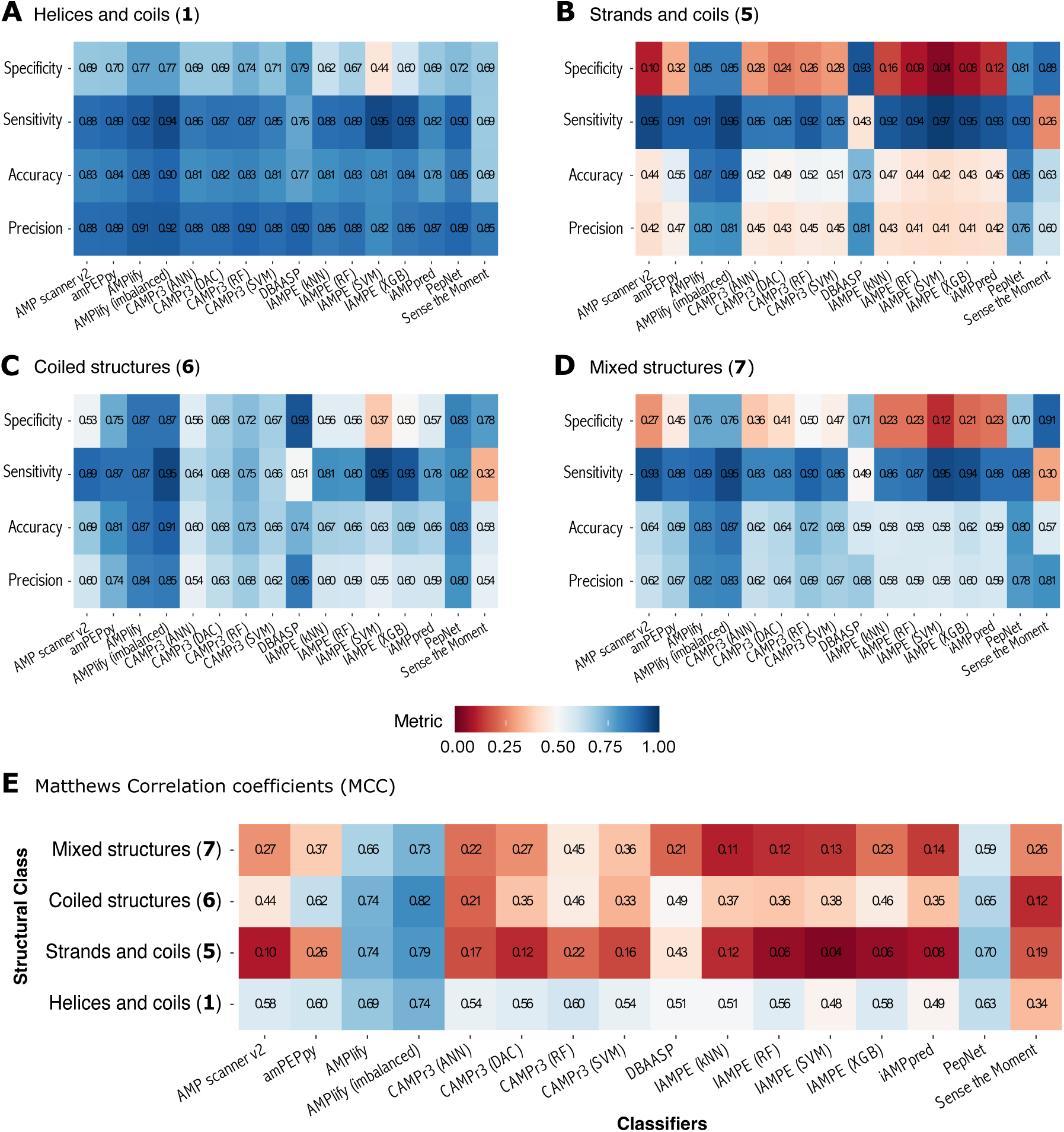
Model performance varies across structural classes: (A) class 1 (helices/coils), (B) class 5 (strands/coils), (C) class 6 (predominantly coiled), and (D) class 7 (mixed). Evaluation is based on confusion-matrix metrics (specificity, sensitivity, accuracy, precision), with global performance captured by Matthews correlation coefficients (E).

For the helical and coil dataset (1, **Figures 2A** and **S1**), most classifiers achieved accuracy and precision consistently above 0.80, indicating robust classification of both AMPs and non-AMPs across all models. Here, the *Sense the Moment* model is a relative underperformer with an accuracy dropping to nearly 0.70. In addition, nearly all models also showed high sensitivity (∼0.80-0.95) but lower specificity (∼0.60-0.80), suggesting a bias toward predicting the positive (AMP) class. As such, the model *IAMPE (SVM)* exhibited the largest gap between sensitivity (∼0.95) and specificity (∼0.42). The models *AMPlify* and *AMPlify (imbalanced)* offer the best balance with relatively high accuracy, precision, and sensitivity without low specificity.

This sensitivity-specificity imbalance persists across other structural classes (5, 7) and classifiers, suggesting a systemic bias in the benchmark and reflecting the composition of the training data. Most predictors in the strands and coils class (5; **Figures 2B** and **S2**) underperformed, with low accuracy, precision (∼0.40), and specificity (∼0.25), but exhibited the highest sensitivity (>0.90). Interestingly, three classifiers *– AMPlify*, *AMPlify (imbalanced),* and *PepNet –* displayed all classification metrics above 0.75, demonstrating strong predictive performance for this structural class. We observed similar trends in the mixed structures class (7, **Figures 2D** and **S4**), where predictors performed moderately well, with accuracy and precision in the 0.60-0.70 range, specificity averaging 0.30, and sensitivity above 0.90. Finally, several classifiers for the coiled class (6, **Figures 2C** and **S3**) achieved near-equivalent performance across all metrics, suggesting either a more tractable structural class or a smaller, less discriminating test set. The *IAMPE (SVM)* and *IAMPE (XGB)* exhibited the largest sensitivity-specificity gaps, indicating a bias toward coiled AMPs. The same three classifiers *– AMPlify*, *AMPlify (imbalanced),* and *PepNet –* outperformed the other models, suggesting they capture features relevant to stranded and coiled peptide datasets that others do not.

Despite these observations, confusion-matrix metrics may overestimate performance on imbalanced datasets^48^. To ensure fairer comparisons across structural classes, we also evaluated performance using the Matthews Correlation Coefficient (MCC)^49^, which balances predictions across both positive and negative classes (see **Materials and Methods**). As shown in **Figure 2E**, MCC values varied across structural classes. The *AMPlify* models are the most robust classifiers across all four structural classes, with *AMPlify (imbalanced)* peaking at 0.82 on coiled structures. The two models maintained strong MCC values even on the hardest classes (5: 0.74/0.79 and 7: 0.66/0.73), where other models collapsed. The model *PepNet* is arguably the third most reliable classifier, with moderate MCC values of 0.70 (strands), 0.65 (coils), 0.63 (helices), and 0.59 (mixed structures). In contrast, the model *Sense the Moment (StM)* exhibited the (near-)lowest MCC values in three of the four structural classes: 0.19 (strands), 0.12 (coils), 0.34 (helices), and 0.26 (mixed structures). Its moderate accuracy (**Figure S2**) on helical peptides may be due to class bias. All *IAMPE* variants and the *iAMPpred* model exhibited near-random MCC values on stranded and mixed structures (5 and 7), confirming the high-sensitivity/low-specificity pattern.

Overall, these results confirm that peptide structural features influence model predictions, with β-sheet-containing classes (5 and 7) consistently performing worse than helical and coiled classes (1 and 6). This pattern suggests an algorithmic bias that hinders the classification of AMPs versus non-AMPs within strand-rich structural space. Structural class 1 (helices and coils) achieved the highest performance across evaluation approaches, confirming reliable classification across models. In contrast, strand-rich classes (5 and 7) exhibited high sensitivity but low specificity, and their low MCC values revealed limited overall discriminatory reliability. This MCC-based evaluation aligns with the confusion matrix analyses (**Figures 2A**-**D** and **S1**-**S4**), and MCC provides a more comprehensive representation of model efficacy than accuracy alone, particularly in skewed datasets where sensitivity-specificity trade-offs can obscure predictive power^48^.

Among the 16 classifiers evaluated, *AMPlify* (both models) and *PepNet* showed the strongest and most consistent performance across all structural classes. Notably, the *AMPlify (imbalanced)* variant achieved the highest single MCC value across the entire benchmark (0.82 on coiled structures) and maintained strong discriminative power on the helical and stranded classes alike, suggesting that accounting for structural class imbalance during model training contributes to its robustness. With the exception of these models, our combined analyses indicate that most classifiers struggle to reliably discriminate AMP from non-AMP sequences containing β-sheet motifs, a limitation that accuracy-based metrics alone would have underrepresented.

Next, we assessed per-sequence model performance by calculating the proportion of sequences correctly classified as GRAMPA or non-GRAMPA across different structural classes (**Figure 3**). The figure displays the predictions – true positives (TP), false negatives (FN), false positives (FP), and true negatives (TN) – for the 16 AMP classifiers tested against the four structural classes (1, 5, 6, and 7). Most models achieve high true-positive rates for GRAMPA peptides (dominant dark blue bars on the left) but also produce significant false positives among non-GRAMPA peptides (light red bars on the right). This further confirms the systemic bias in training toward known AMPs. The right side of each panel shows that a few models, such as *AMPlify* and *PepNet*, consistently achieve high true-negative rates (dark red bars) across all four structural classes.

**Figure 3.**
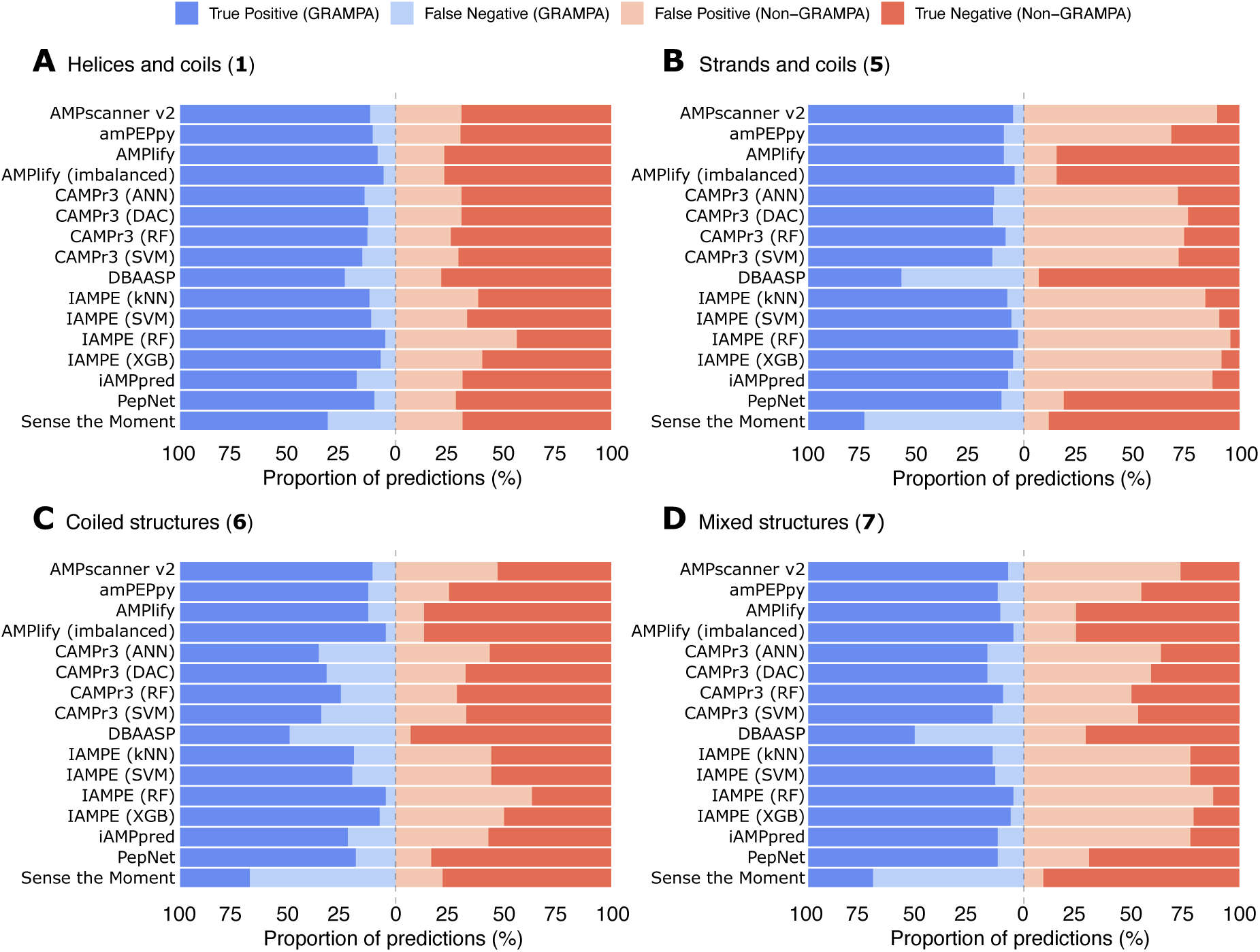
Mapping the proportions of predictions across classifiers and structural classes - (A) 1: Helices and coils, (B) 5: Strands and coils, (C) 6: Mostly coiled structures, and (D) 7: Mixed structures.

In **Figure 3A**, models perform confidently overall against the helical structural class (1), as expected, since *α*-helical peptides are likely to dominate AMP training databases, such as GRAMPA^32^. Most classifiers output high TP rates (about 75-95%) and moderate TN rates for non-GRAMPA peptides. *IAMPE (RF)* has the lowest TN rate; the model may struggle with helical non-AMPs. *Sense the Moment (StM)* performs well on helices, consistent with its design to distinguish AMP sequences from randomized variants that maintain the hydrophobic moment but lack sequence order and secondary structure. The authors of *StM* recommend using their tool as a complementary model, primarily for sequences prone to form helical motifs^45^. The model performance declines in other structural classes.

**Figure 3C** displays greater variation across models, with performance differences falling between those of the other structure classes. Several classifiers show low TP rates (<60%) and high false-negative rates for GRAMPA peptides, especially *DBAASP* and *StM*, indicating that coiled structures are often misclassified as non-AMPs – likely due to underrepresentation or structural ambiguity in the training sets. Among the *IAMPE* variants, differences in performance on coiled structures appear to be driven more by model architecture than by training data, since all variants use the same training set. *IAMPE (RF)* achieves a high TP rate for GRAMPA coils but has the lowest true-negative rate among *IAMPE variants*, suggesting that the random forest architecture tends to overfit AMPs for this structural class. The other *IAMPE* models (kNN, SVM, XGB) attain higher TN rates despite lower TP rates, indicating that these architectures are more conservative in predicting coiled structures. A similar pattern is observed among *CAMPr3* variants, with the random forest model achieving the highest TP and TN rates in the series. *AMPlify*, *AMPlify (imbalanced), amPEPpy*, and *PepNet* performed best on this structural class, achieving the highest combined true-positive and true-negative rates.

In **Figures 3B** and **3D**, overall performance drops notably, reflecting that stranded AMPs (class 5) and mixed structures (class 7) are underrepresented in the training data. Many models – including *AMP scanner v2*, *CAMPr3* variants, and *IAMPE* variants – misclassify non-GRAMPA strands and mixed structures as AMPs, resulting in higher false-positive rates despite maintaining high TP rates for GRAMPA sequences. This simultaneous rise in both true-positive and false-positive rates suggests that these models recognize broad physicochemical features common to helical AMPs rather than structure-specific signatures. Conversely, *DBAASP* and *StM* show the opposite pattern – with higher false-negative and true-negative rates – consistent with a training bias toward helical AMPs and, in the case of *StM*, its original helix-centric design. Among all tested models, *AMPlify*, *AMPlify (imbalanced)*, and *PepNet* perform best across these two structural classes, exhibiting relatively higher TP and TN rates, which aligns with their strong performance shown in **Figure 3C**.

To better understand model confidence, we examined class probability distributions for nine out of the 16 classifiers that provided probability outputs (**Figure S5**). Overall, GRAMPA peptides (shown in blue) had median probabilities above 0.5 across all structural subsets, consistent with their correct labelling as AMPs. In contrast, non-GRAMPA sequences (depicted in red) exhibited wider distributions and greater uncertainty. Structural classes lacking stranded motifs (classes 1 and 6) typically fell below the 0.5 threshold, while strand-rich subsets (classes 5 and 7) often exceeded it—consistent with the higher false positive rates seen in **Figure 3**. Model confidence also mirrored structural bias: probability distributions are tighter and higher for helical GRAMPA peptides, gradually broadening across coiled, stranded, and mixed structural classes, reinforcing the training bias shown in **Figure 3**. Among the nine models, *AMPlify* and *AMPlify (imbalanced)* produce the clearest separation between GRAMPA and non-GRAMPA probability distributions across all structural classes, with tight distributions approaching 1.00 for GRAMPA sequences and 0.00 for non-GRAMPA sequences. *PepNet* displayed similarly well-separated distributions for most structural classes, though notably more dispersed probabilities for non-GRAMPA helices and mixed structures. In contrast, non-GRAMPA sequences with stranded or mixed structures were assigned intermediate class probabilities (0.25-0.75) across most remaining models - including *amPEPpy*, *AMP scanner v2*, *CAMPr3* variants, and *iAMPpred* - indicating that their decision boundaries are poorly calibrated for non-helical structures.

In summary, structural class strongly influences classifier performance, with helical peptides being the most consistently detected and β-sheet-rich subsets (5 and 7) presenting the greatest challenge. Most models exhibit a high false-positive rate across structural classes, reflecting limited specificity for non-helical AMPs and suggesting that current classifiers generalize poorly beyond the helical sequences that dominate their training data. Few models perform consistently across all four structural classes; *AMPlify*, *AMPlify (imbalanced)*, and *PepNet* demonstrated the most balanced profiles in **Figure 3**, and the probability distributions in **Figure S5** further corroborate their superior discrimination across all structural classes.

### 3.3. Understanding structural bias in predictions

To elucidate the origins of observed prediction biases, particularly the misclassification of non-GRAMPA sequences containing β-sheet motifs as AMPs, we investigated two potential contributing factors: structural imbalance and data leakage in training sets.

We first assessed whether structural diversity was adequately represented across both sequence classes. We encoded the structures of both external GRAMPA and non-GRAMPA validation sets using ProstT5^46^, a sequence-structure encoder that generates 3Di tokens as computationally efficient proxies for three-dimensional representations. The approach circumvented the need for explicit structure predictors such as PEP2D and AlphaFold2. The resulting structural embeddings were projected onto a two-dimensional UMAP^47^ manifold fine-tuned to maximize the separation of the four structural classes under consideration (folds 1, 5, 6, 7) – see **Figure S6**. As shown in **Figure 4**, the two datasets are distributed across the structural space, forming two clusters: a broad, continuous lower manifold (bottom center-right) and a compact upper cluster (top left). The first cluster contains the four predicted folds, predominantly dominated by structural classes 1 and 6, whereas the second cluster includes only structural classes 5 and 7. Comparing the datasets fold by fold, GRAMPA helices assigned to fold 1 (light green, panel A) are abundant, whereas their non-GRAMPA counterparts (yellow, panel B) remain underrepresented. Conversely, non-GRAMPA sequences are enriched in strands and coils (5 and 6, red and orange) relative to their GRAMPA counterparts. These asymmetries mirror the structural imbalance documented in **Table 1**, namely, the overrepresentation of helical folds among AMPs and of strands and coils among non-AMPs – a bias that likely underlies the misclassification of stranded and coiled non-GRAMPA sequences as AMPs.

**Figure 4.**
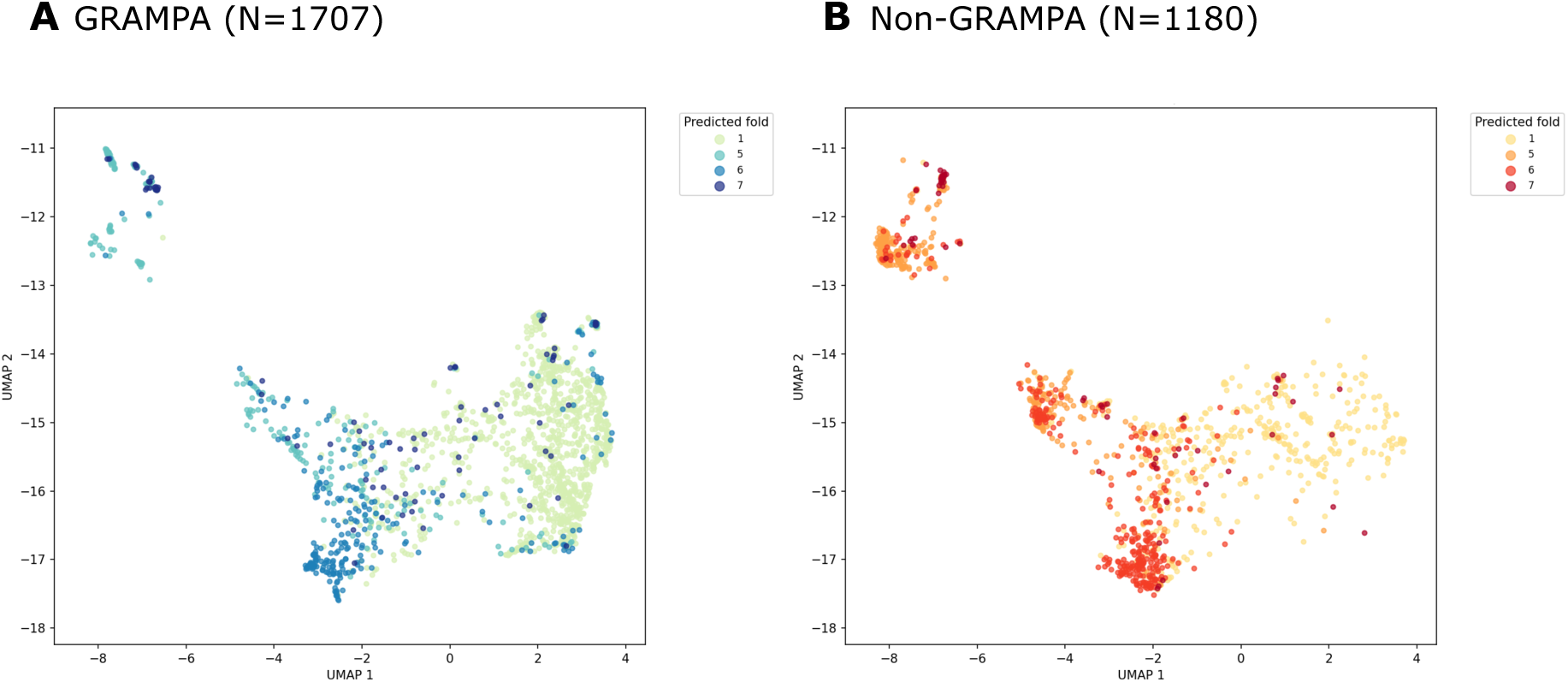
Structural diversity of GRAMPA and non-GRAMPA external validation sets projected onto a UMAP structural space. UMAP projections of ProstT5-derived 3Di structural embeddings for the GRAMPA (A) and non-GRAMPA (B) validation sets, colored by predicted structural fold (1, 5, 6, 7). The manifold was fine-tuned to maximize the separation of the four structural classes.

We then compared the structural distributions of the two external validation sets with those of the available training sets for four classifiers: GRAMPA against the positive training sets and non-GRAMPA against the negative training sets. The classifiers were *AMP scanner v2* (panels A and B), *amPEPpy* (panels C and D), *AMPlify*, and *PepNet* sharing identical training sets (panels E and F). The results are illustrated in **Figure 5**.

**Figure 5.**
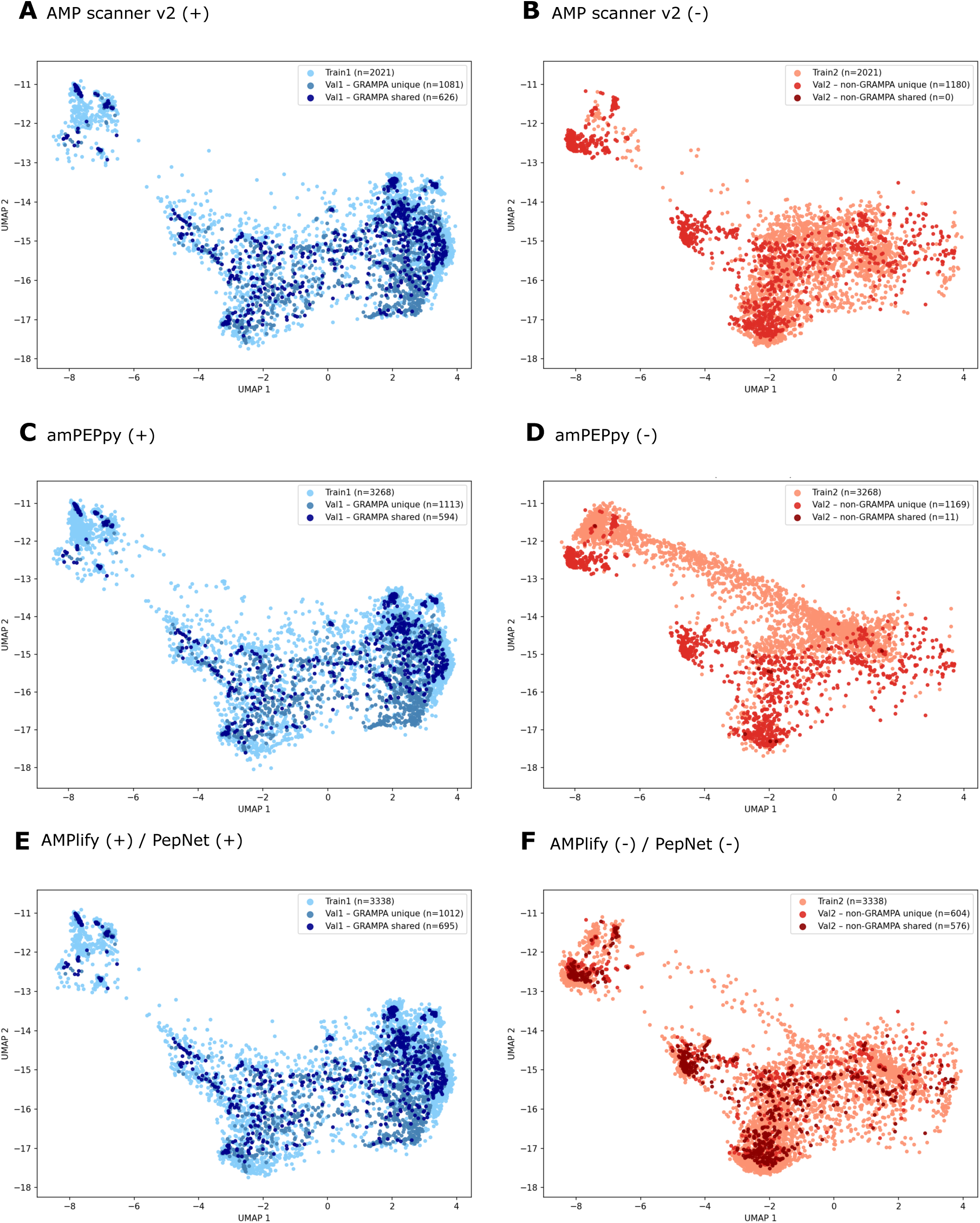
Structural overlap between external validation sets and classifier training sets in UMAP space. UMAP projections of ProstT5-derived 3Di structural embeddings compare the structural distributions of GRAMPA (Val1, blue shades) and non-GRAMPA (Val2, red shades) with the positive and negative training sets (light shades) for *AMP scanner v2* (A, B), *amPEPpy* (C, D), and *AMPlify/PepNet* (E, F), respectively. Darker points denote the validation sequences shared with the corresponding training set. Sample sizes are indicated in the legends.

Across all four classifiers, a large fraction of GRAMPA sequences overlaps with the positive training sets (dark blue, “shared”), indicating varying degrees of data leakage (defined here as sequence identity) into them. The shared sequences are not confined to a specific structural region but are distributed broadly across the UMAP manifold, suggesting the leakage is uniform across structural classes. The proportion of shared sequences is notable: 626/1707 for *AMP scanner v2*, 594/1707 for *amPEPpy*, and 695/1707 for *AMPlify/PepNet*, representing roughly one third of GRAMPA sequences are shared with the respective positive training sets. Of note, *AMPlify (imbalanced)* shares the same positive training set as AMPlify and PepNet and therefore exhibits an identical positive leakage (695/1707).

On the negative side, *AMP scanner v2* (panel B) and *amPEPpy* (panel D) are effectively leakage-free; both show zero and 11 shared sequences, respectively, between the non-GRAMPA validation set and the negative training sets. *AMPlify/PepNet* (panel F) includes 576 shared sequences out of 1180, representing nearly half of the non-GRAMPA validation set that overlaps with the negative training data. *AMPlify (imbalanced)* retains this negative leakage and expands to over 102,000 peptide sequences (not depicted). Consequently, the fraction of non-GRAMPA sequences is at least as high as in panel F, increasing the likelihood that additional non-GRAMPA sequences are covered.

The models *AMP scanner v2* and *amPEPpy* exhibit negligible negative leakage but perform poorly on non-GRAMPA strands and mixed structures (5 and 7; **Figures 3B** and **3D**). The negative training sets (light red) are heavily concentrated in the lower broad cluster, dominated by helices and coils (1 and 6). The non-GRAMPA validation sequences (dark red, “unique”), however, are distributed across both the lower manifold and the upper cluster, which is occupied by folds 5 and 7 (strands and mixed structures). *AMP scanner v2* (panel B) contained no negative stranded and mixed examples, whereas *amPEPpy* (panel D) spans a broader, more continuous structural space, partially covering the upper cluster with fewer helical folds (1) - yet this partial coverage remains insufficient to prevent misclassification of stranded and mixed structures. This structural underrepresentation in the negative training sets causes both models to misclassify out-of-distribution non-GRAMPA sequences from folds 5 and 7 as AMPs by default.

The broader negative set coverage visible in panel F (**Figure 5**) offers a structural explanation for the apparent superior performance of *AMPlify/PepNet* across structural classes, including greater representation of stranded and mixed structures in the upper cluster. However, the data leakage observed in panels E (∼41% of GRAMPA) and F (∼49% of non-GRAMPA) indicates that both validation sets are contaminated with sequences from the *AMPlify, AMPlify (imbalanced)*, *PepNet* training datasets. All three models may simply be recognizing examples seen during training (memorization) rather than learning structurally generalizable discriminating features. In the case of *AMPlify (imbalanced)*, the disproportionate size of the negative training set increases the likelihood that non-GRAMPA validation sequences are represented in training. Their superior performance across all four structural classes may therefore be inflated by this leakage, making it difficult to determine how much is attributable to learning versus memorization of identical training examples.

To disentangle data leakage from genuine pan-fold learning, we re-evaluated classification performance on non-overlapping subsets of GRAMPA (N=1,012) and non-GRAMPA sequences (N=604). Sustained superior discrimination on these subsets would support genuine model quality, whereas a performance drop would implicate data leakage as a major confounding factor. We report prediction results – true positives (TP), false negatives (FN), false positives (FP), and true negatives (TN) – for the *AMPlify* and *PepNet* models, both before (**Tables S2** and **S4**) and after removing duplicate sequences (**Tables S3** and **S5**). These results are summarized in **Figure 6**.

**Figure 6.**
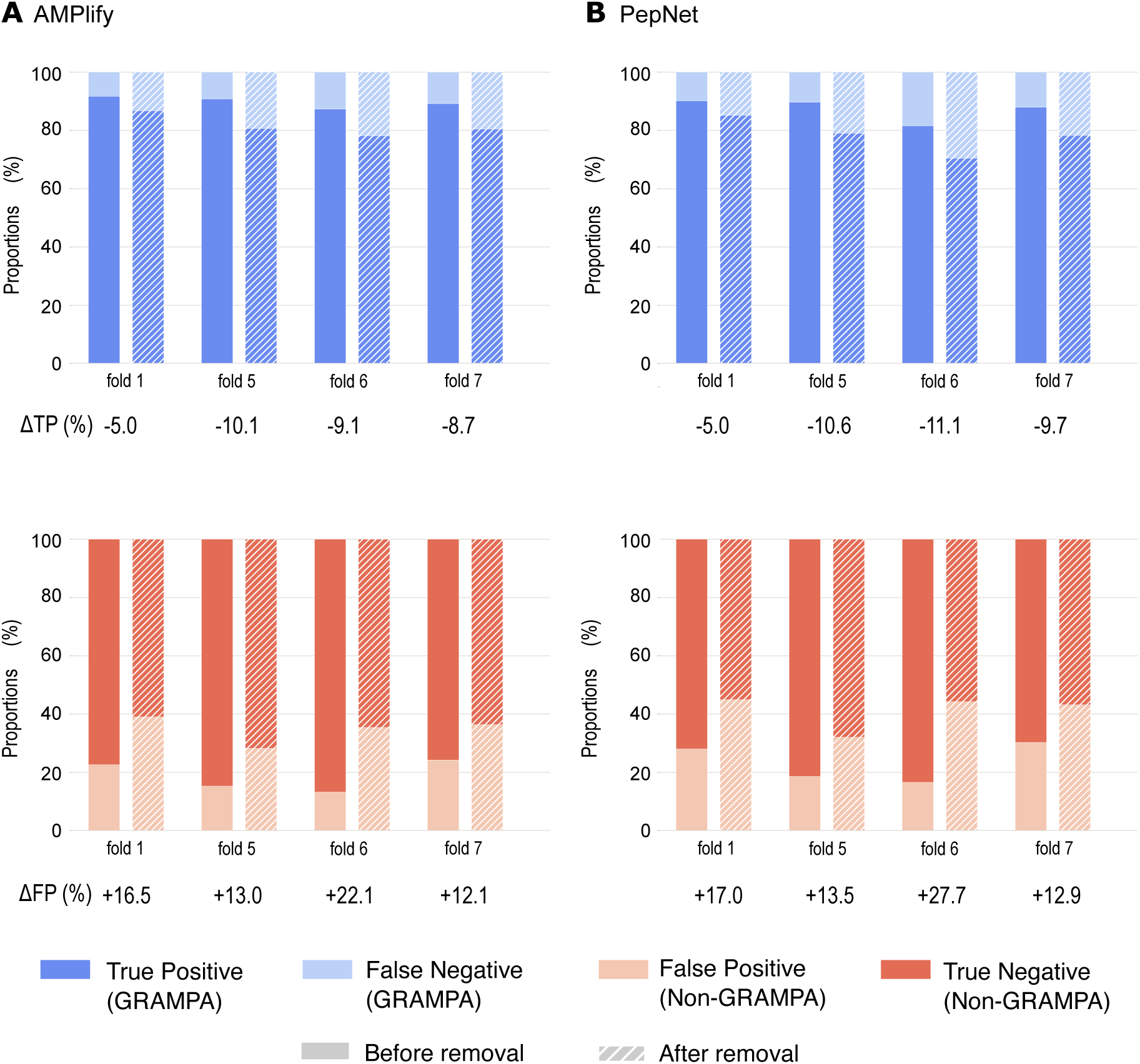
Classification proportions before and after data leakage removal for AMPlify (A) and PepNet (B). The positive set (GRAMPA/AMP) is shown in the upper panels as stacked proportions of TP (dark blue) and FN (light blue). The negative set (non-GRAMPA/non-AMP) is shown in the lower panels as stacked proportions of FP (light red) and TN (dark red). Solid bars represent performance before leakage removal; hatched bars represent performance after removing shared sequences between training and validation sets. Delta values (ΔTP%, ΔFP%) indicate the change in proportion after leakage removal. Folds correspond to structural classes: 1 (helices), 5 (strands), 6 (coils), and 7 (mixed structures).

The removal of shared sequences between the training and validation sets resulted in significant losses for both GRAMPA and non-GRAMPA sequences across folds. GRAMPA lost 37.4% of helices (fold 1), half (52.5%) of stranded AMPs (fold 5), 41.8% coils (fold 6), and 44.6% mixed structures (fold 7). Similarly, non-GRAMPA diminished 44% of helices, 46% of stranded non-AMPs, nearly two-thirds (62.6%) of coils, and one-third of mixed structures.

After removing leakage, *AMPlify* consistently outperforms *PepNet* across all folds in both sensitivity and specificity (**Figure 6**). For *AMPlify*, FN counts are exactly preserved across all folds, and FP counts are nearly identical (fold 1: 96 to 93), indicating that the performance drops are almost entirely denominator-driven; consistent with the near-complete memorization of the leaked sequences. For *PepNet*, both FN and FP counts decrease slightly after de-leaking, indicating that a small number of leaked sequences were misclassified and that memorization was therefore less complete than in *AMPlify*. Sensitivity (ΔTP%) drops 5-11% for both models across folds, while the false positive rate (ΔFP%) rises by 13-22% for *AMPlify* (**Figure 6A**) and 13-28% for *PepNet* (**Figure 6B**). Specificity is more affected than sensitivity across all folds and both models, suggesting that the non-AMP decision boundary has learned less from genuine sequence features and is more vulnerable to leakage removal.

Examining performance by structural class reveals consistent patterns. Fold 1 (helices) retains the highest sensitivity for both models (*AMPlify* 86.7%, *PepNet* 84.8%) and shows the smallest sensitivity drop (ΔTP% = –5.0% for both), suggesting that helical AMPs carry more discriminative sequence features beyond the training set. Fold 5 (stranded AMPs – upper panels) exhibits greater sensitivity loss relative to baseline (ΔTP% = –10.1% and –10.6%), consistent with the largest proportion of leaked sequences (52.5%). Fold 7 (mixed structures) shows more modest declines in either sensitivity or specificity across both models, despite its small sample size. The largest divergence between models occurs in fold 6 (coiled structures), where both models fall below 75% sensitivity. With *PepNet*, the proportion of true positives decreases from 81.6% to 70.4% after leakage removal (**Tables S4** and **S5**). *AMPlify* retains approximately 8% higher sensitivity and 9% lower FP rate than *PepNet*. Overall, these results indicate that both models had inflated performance metrics due to data leakage, with coiled AMPs representing a structurally distinct class that neither model has learned to reliably distinguish from non-AMPs on novel sequences.

## 4. DISCUSSION

Our study demonstrates that current antimicrobial peptide (AMP) predictive models exhibit structural bias arising from the uneven representation of peptide structural classes in training datasets. Most models achieve high predictive accuracy for sequences adopting helical folds and coils, but perform poorly on β-sheet motifs, thereby limiting their applicability across the broader AMP structural space. These observations are consistent with independent work by Dean and co-workers, who reported that computationally predicted and generated AMPs against *E. coli*, *S. aureus,* and *P. aeruginosa* predominantly folded into α-helices^50^. This mirrors the inherent bias towards helical motifs in existing AMP datasets used for activity prediction and generation. Comparable biases have been reported across other protein classes^51^ and computational tasks, resulting in skewed predictions in functional annotation^52,53^, protein-protein interaction networks^27^, protein stability^54,55^, protein structure^32,56^, and in sequence or structure generation^57,58^.

This work contributes to a growing body of studies showing that biases in training data inflate performance metrics in protein prediction and generation. Multiple strategies have been proposed to mitigate systematic bias in ML models, including class resampling, clustering-based methods, synthetic data augmentation *via* generative algorithms, feature augmentation, and optimized classification algorithms^55,59^. In our recent work, subset selection enabled the development of structure-specific models with superior predictive performance, whereas targeted data reduction led to information loss in structure-agnostic models^60^, underscoring the need for structural diversity and rigorous curation in the next generation of ML models^56^.

The structural biases documented here also carry underappreciated consequences for prospective experimental campaigns. Poor model recall of β-sheet AMPs leads to fewer stranded candidates being synthesized and tested, fewer representatives in public databases, and continued training imbalance in future models – a self-reinforcing loop analogous to chemotype bias in drug discovery^61–63^. Breaking this cycle requires treating structural diversity as an explicit experimental design criterion, prioritizing β-sheet and mixed-fold candidates for synthesis over relying solely on predicted activity scores.

The observed differences in classification performance across structural classes are attributable to training imbalance and variation in learned (physicochemical) features across folds. α-Helical AMPs possess well-characterized cationic charge distribution, hydrophobicity, and amphipathicity that are captured by the hydrophobic moment, a vector sum of side-chain hydrophobicities projected along the helical axis and the basis of the *StM* model^45,64^. These properties are amenable to extraction by both traditional machine learning and neural network architectures. By comparison, stranded and mixed AMPs include structural components (e.g., hydrogen bonding, strand orientation, disulfide bridges) that are non-local and poorly captured by sequence-only encodings. The superior sensitivity retained for helical AMPs after leakage removal (fold 1: *AMPlify* 86.7%, *PepNet* 84.8%) reflects the stronger discrimination in sequence space for this class and points to an intrinsic limitation of sequence-only models for stranded and coiled structures, motivating the development of structure-aware feature representations for pan-fold discrimination of AMPs.

Beyond the structural imbalance, the de-leaking analysis further reveals that the two top-performing models exhibited inflated performance metrics driven by near-complete memorization of the leaked sequences rather than generalization. Some parallels can be drawn with large generative models, where training on insufficiently diverse data may lead to memorization that masks poor generalization. The accompanying sensitivity loss (5-11%) and increase in false positive rate (13-28%) highlight that neither model has learned to reliably distinguish non-AMPs from novel sequences in the coiled structural space. A principled mitigation strategy is to develop selective classifiers that abstain from making predictions when an input falls outside the structural training distribution^65^, for which the ProstT5-UMAP projections reported here provide a natural basis.

Finally, our benchmarking analysis compared models of increasing algorithmic complexity: early models (pre-2022) primarily relied on SVM and Random Forest classifiers, while recent models incorporate CNNs or transformers. Greater complexity has improved predictive performance but has not resolved the underlying sampling bias. Joint sequence-structure embeddings (e.g., Prost T5^46^, Saprot^66^) and co-generation approaches that pair sequence and structure data are promising avenues, though they currently risk biasing exploration toward well-represented folds. As ProstT5 was trained on AlphaFold2 predictions that were filtered for high structural confidence, enriched in well-structured helical proteins, and excluded disordered sequences, the derived embeddings should be treated with caution. With sufficiently diverse and balanced training data, these approaches hold the potential to expand exploration across broader protein structural space^67^.

## 5. CONCLUSION

Our study highlights structural bias in predictive models for identifying novel antimicrobial peptides. The combination of uneven structural representation in training data, data leakage, and the intrinsic learnability of helical versus non-helical folds limits the generalization and reliability of current AMP classifiers across the full structural space. While current efforts to integrate sequence, structure, and function into machine learning-guided frameworks offer more controllable peptide and protein design with improved success rates, they may also constrain novelty by reinforcing exploration of familiar structural territory. Addressing structural bias – through curated, structurally diverse training data, selective prediction strategies, and joint sequence-structure modeling – will be essential to unlocking the ‘dark’ peptidome and proteome, creating opportunities for novel discoveries that could drive the next generation of therapeutics, biopesticides, and biomaterials.

## Data and code availability

Datasets include peptide sequences, ProsT5 embeddings, activity, and fold predictions. R and Python scripts to reproduce ternary plots and UMAP-ProsT5 projections are all available at https://github.com/plissonf/AMP-structural-bias-audit.

## Declaration of competing interest

V.D.A.-B. declares no competing interests. F.P. is the founder and scientific director of Ingenie Bio, a company that provides consulting and services in computational peptide design.

## Acknowledgments

The authors thank the Mexican *Secretaría de Ciencias, Humanidades, Tecnología e Innovación* (SECIHTI, formerly CONAHCYT), for financial support through grant A1-S-32579 (2019-2023) supporting this study. V.D.A.-B. acknowledges a national postgraduate scholarship awarded by CONAHCYT during the funded period.

## Author contributions

F.P. conceptualized the investigation. V.D.A.-B. and F.P. carried out the investigation, including methodology, data curation, benchmarking, and bioinformatics analysis. Both authors wrote, edited, and reviewed the manuscript.

## Notes

https://github.com/plissonf/AMP-structural-bias-audit

