## Supporting Information for "Structural bias in machine learning-guided peptide design"

### Supplementary Information

**Table S1**. Binary classifiers predicting antimicrobial activity. 2

**Table S2**. Proportions of *AMPlify* predictions on GRAMPA (N=1,707) and non-GRAMPA (N=1,180) validation sets and across folds. 9

**Table S3**. Proportions of *AMPlify* predictions on GRAMPA (N=1,012) and non-GRAMPA (N=604) validation sets and across folds – leakage-free. 9

**Table S4**. Proportions of *PepNet* predictions on GRAMPA (N=1,707) and non-GRAMPA (N=1,180) validation sets and across folds. 9

**Table S5**. Proportions of *PepNet* predictions on GRAMPA (N=1,012) and non-GRAMPA (N=604) validation sets and across folds – leakage-free. 9

**Figures S1.** AMP classifier performance on helical peptides (1). 3

**Figures S2.** AMP classifier performance on stranded peptides (5). 4

**Figures S3.** AMP classifier performance on coiled structures (6). 5

**Figures S4.** AMP classifier performance on mixed structures (7). 6

**Figure S5.** AMP class probability distributions across folds. 7

**Figure S6.** UMAP hyperparameter optimization landscape. 8

**Table S1**. Binary classifiers predicting antimicrobial activity.

| Year | Classifier | Algorithm | URL | # |
|---|---|---|---|---|
| 2016 | **CAMPR3** | Support Vector Machine | http://www.camp.bicnirrh.res.in | 1 |
| | | Artificial Neural Network | | |
| | | Random Forest | | |
| | | Discriminant Analysis | | |
| 2017 | **iAMPpred** | Support Vector Machine | http://cabgrid.res.in:8080/amppred/ | 2 |
| 2018 | **AMP scanner V2** | Deep Neural Network | https://www.dveltri.com/ascan/ | 3 |
| 2020 | **IAMPE** | Support Vector Machine | http://cbb1.ut.ac.ir/AMPClassifier/ | 4 |
| | | Random Forest | | |
| | | *k*-Nearest Neighbor | | |
| | | eXtreme Gradient Boosting | | |
| 2021 | **amPEPpy** | Random Forest | https://github.com/tlawrence3/amPEPpy | 5 |
| 2022 | **Sense the Moment (StM)** | Hydrophobic moment threshold | https://www.portoreports.com/stm | 6 |
| 2022 | **DBAASP** | Physicochemical properties thresholds | https://dbaasp.org/ | 7 |
| 2023 | **AMPlify** | Deep neural network including bi-LSTM, MHSDPA and CA layers | https://github.com/bcgsc/AMPlify | 8 |
| 2024 | **PepNet** | Pre-trained protein language model | http://liulab.top/PepNet | 9 |

**Figures S1. AMP classifier performance on helical peptides (1).** Bar chart comparing accuracy (yellow), precision (blue), sensitivity (red), and specificity (green) for 16 AMP classifiers evaluated on a benchmark dataset of helical peptides.

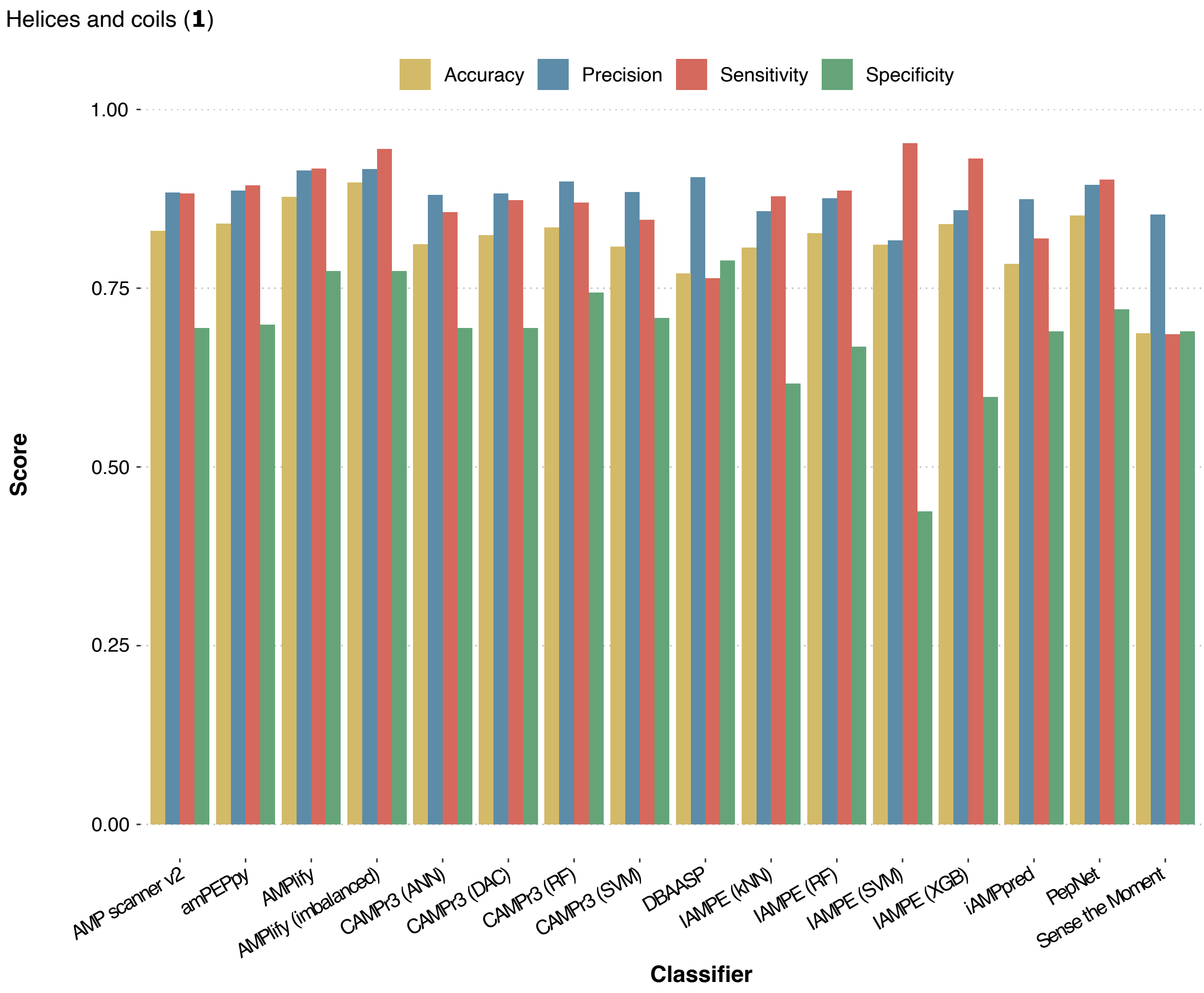

**Figures S2. AMP classifier performance on stranded peptides (5).** Bar chart comparing accuracy (yellow), precision (blue), sensitivity (red), and specificity (green) for 16 AMP classifiers evaluated on a benchmark dataset of stranded peptides.

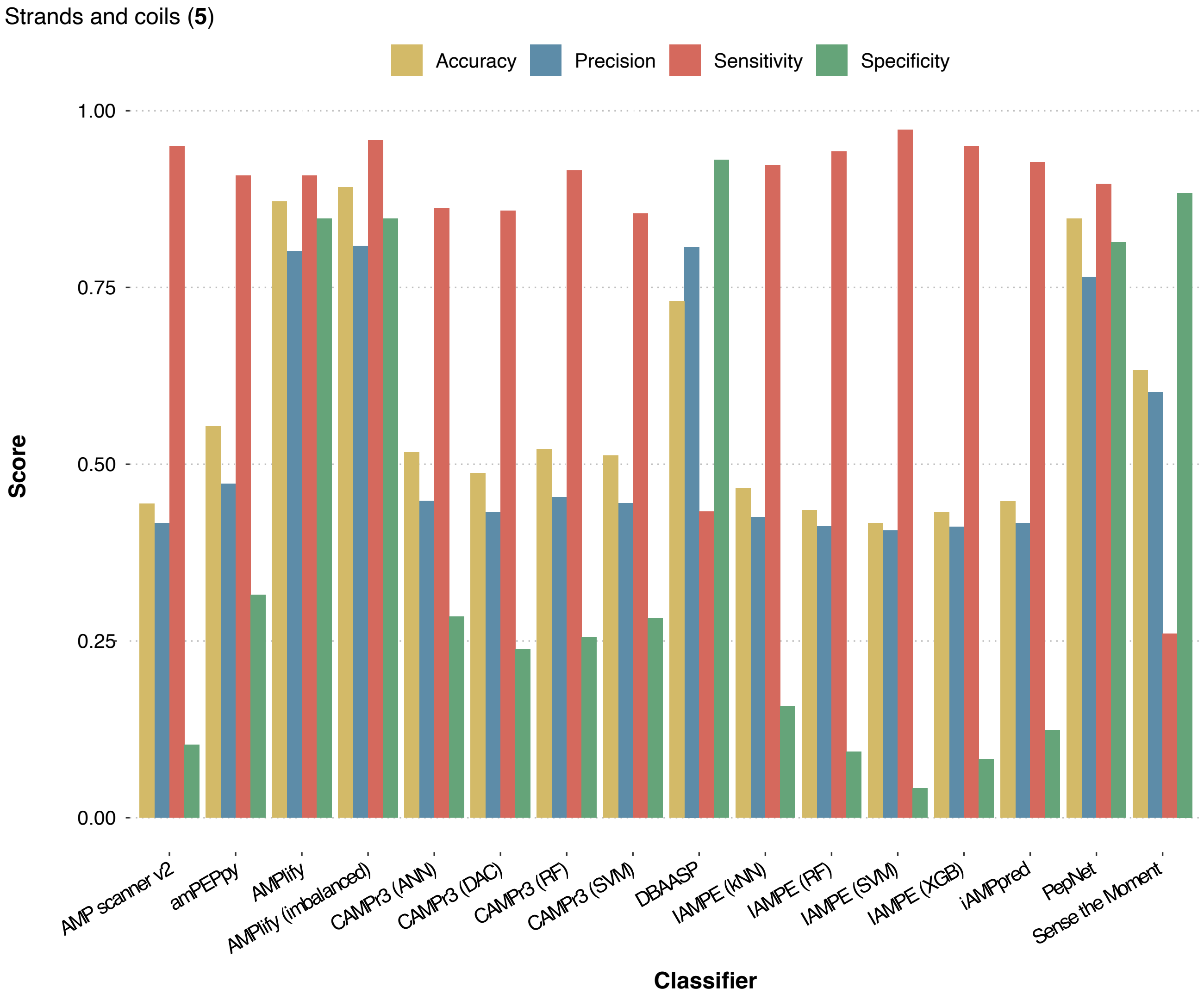

**Figures S3. AMP classifier performance on coiled structures (6).** Bar chart comparing accuracy (yellow), precision (blue), sensitivity (red), and specificity (green) for 16 AMP classifiers evaluated on a benchmark dataset of coiled peptides.

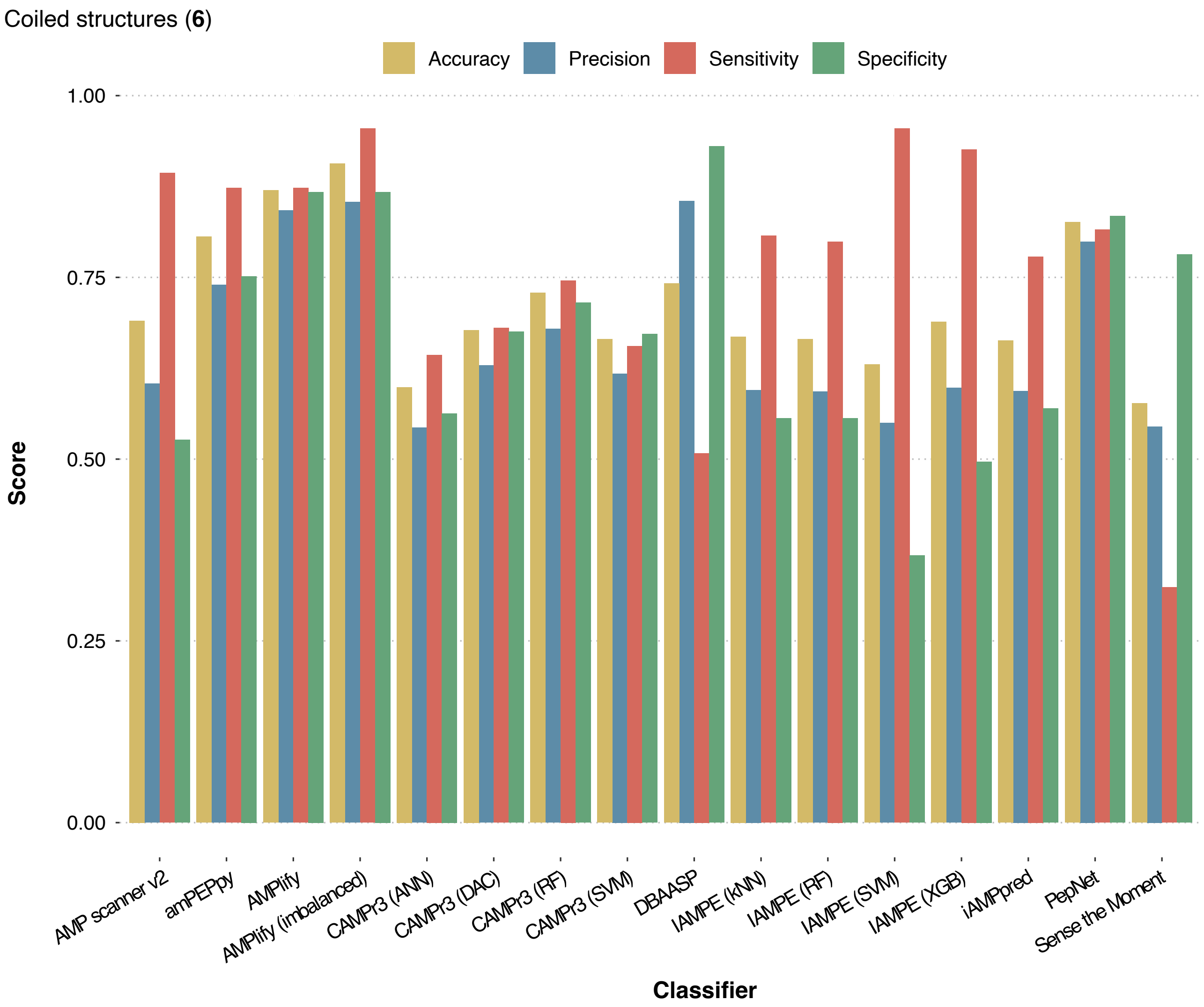

**Figures S4. AMP classifier performance on mixed structures (7).** Bar chart comparing accuracy (yellow), precision (blue), sensitivity (red), and specificity (green) for 16 AMP classifiers evaluated on a benchmark dataset of peptides folding into mixed structures.

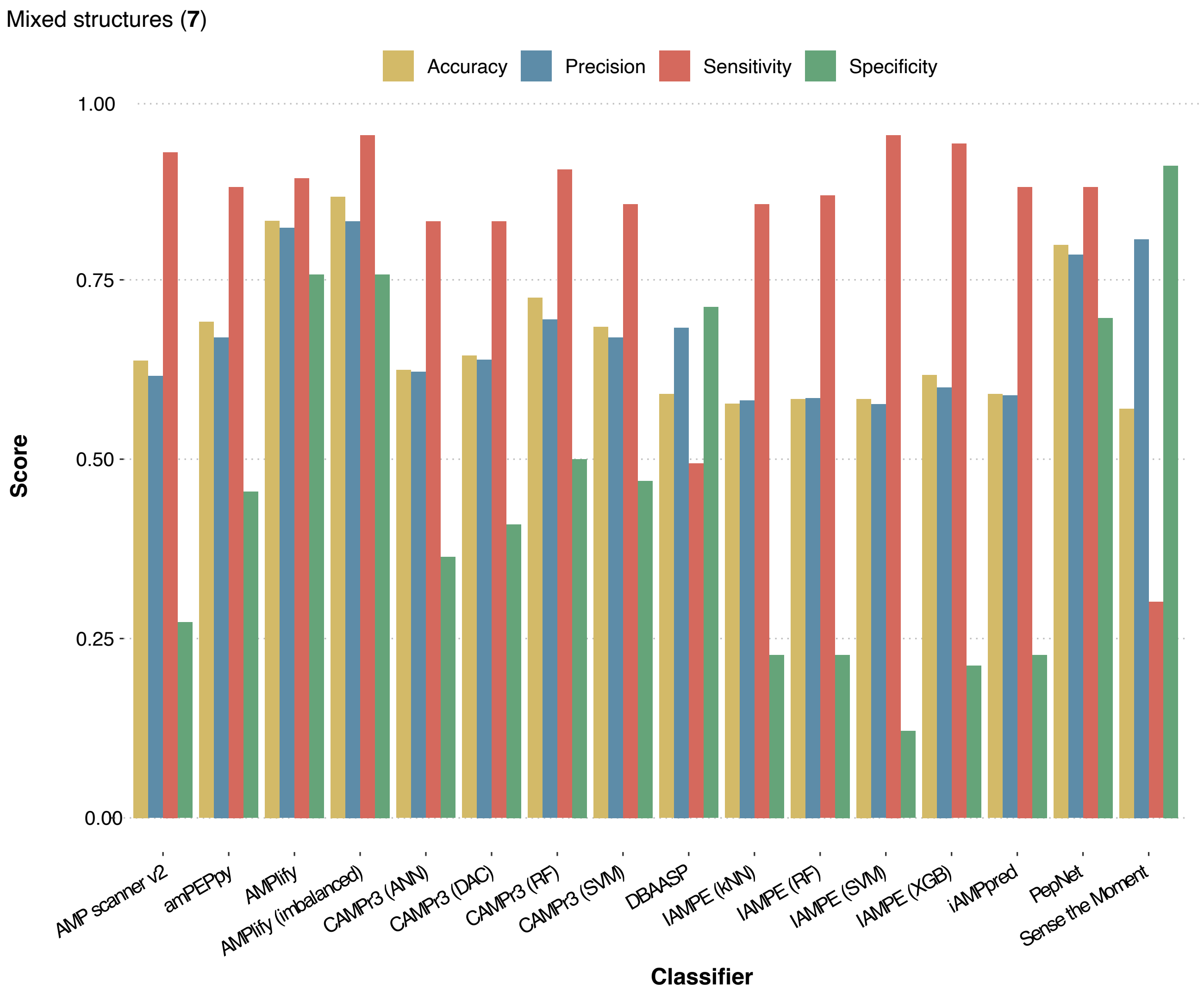

**Figure S5. AMP class probability distributions across folds**. Box plots show the distributions of AMP class probabilities assigned by nine classifiers (*amPEPpy*, *AMP scanner v2*, *CAMPr3* [*DAC*, *RF*, *SVM*], *iAMPpred*, *AMPlify* [*balanced*, *imbalanced*], and *PepNet*) to the GRAMPA (blue) and non-GRAMPA validation sets across the four structural classes, helices, coils, strands, and mixed structures.

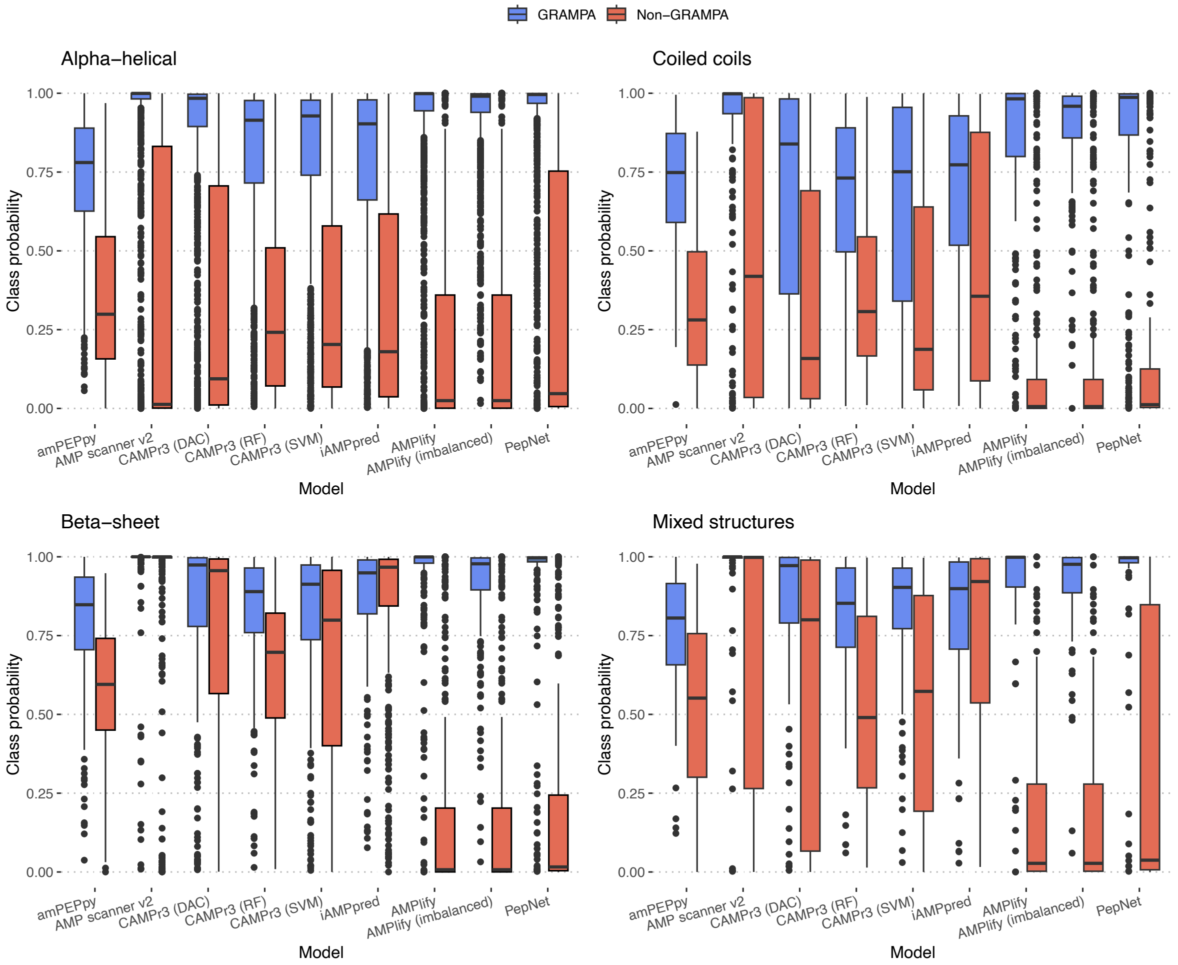

**Figure S6. UMAP hyperparameter optimization landscape.** Heatmap of silhouette scores across *n_neighbors* (y-axis, 25-2886) and *min_dist* (x-axis, 0.00-0.50) for three distance metrics (cosine, correlation, euclidean). Color intensity encodes *silhouette score* on a shared global scale (light = low, dark teal = high). Dashes denote parameter combinations not evaluated.

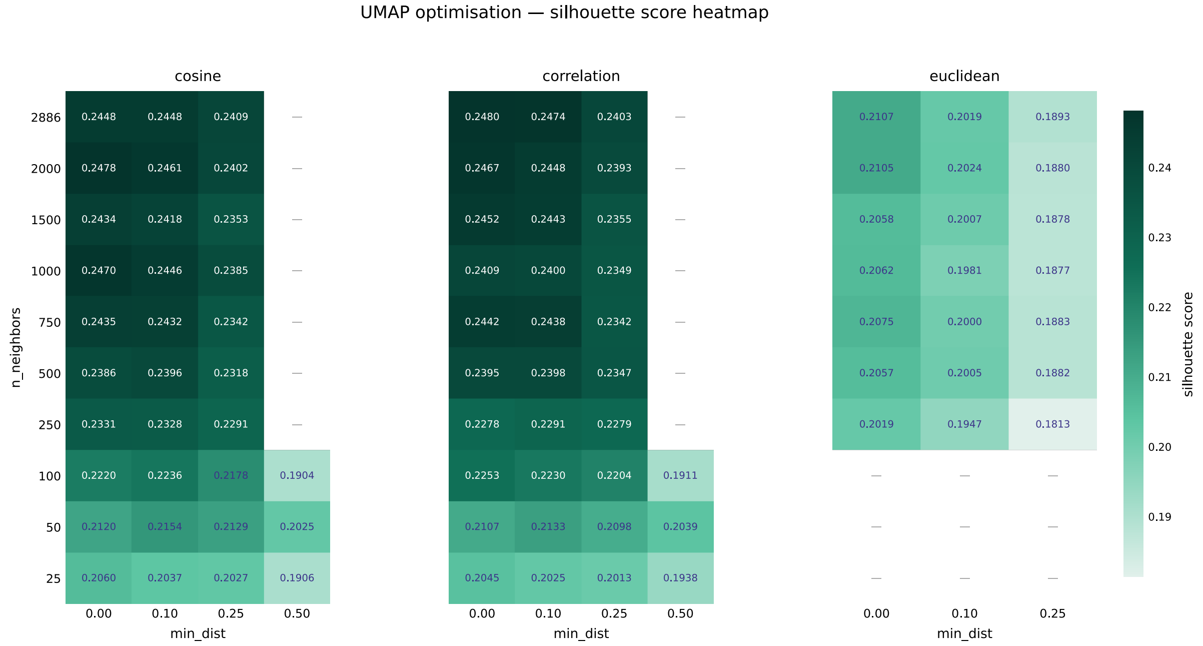


For comparison, the PCA silhouette score is 0.1956 (PC1=0.297, PC2=0.153).

**Table S2**. Proportions of *AMPlify* predictions on GRAMPA (N=1,707) and non-GRAMPA (N=1,180) validation sets and across folds.

| | GRAMPA | | | | | Non-GRAMPA | | | | |
|---|---|---|---|---|---|---|---|---|---|---|
| *Fold* | *Total* | *TP* | *(%)* | *FN* | *(%)* | *FP* | *(%)* | *TN* | *(%)* | *Total* |
| 1 | 1119 | 1026 | (91.7) | 93 | (8.3) | 96 | (22.6) | 329 | (77.4) | 425 |
| 5 | 261 | 237 | (90.8) | 24 | (9.2) | 59 | (15.2) | 328 | (84.8) | 387 |
| 6 | 244 | 213 | (87.3) | 31 | (12.7) | 40 | (13.2) | 262 | (86.8) | 302 |
| 7 | 83 | 74 | (89.2) | 9 | (10.8) | 16 | (24.2) | 50 | (75.8) | 66 |

**Table S3**. Proportions of *AMPlify* predictions on GRAMPA (N=1,012) and non-GRAMPA (N=604) validation sets and across folds – leakage-free.

| | GRAMPA | | | | | Non-GRAMPA | | | | |
|---|---|---|---|---|---|---|---|---|---|---|
| *Fold* | *Total* | *TP* | *(%)* | *FN* | *(%)* | *FP* | *(%)* | *TN* | *(%)* | *Total* |
| 1 | 700 | 607 | (86.7) | 93 | (13.3) | 93 | (39.1) | 145 | (60.9) | 238 |
| 5 | 124 | 100 | (80.6) | 24 | (19.3) | 59 | (28.2) | 150 | (71.8) | 209 |
| 6 | 142 | 111 | (78.2) | 31 | (21.8) | 40 | (35.4) | 73 | (64.6) | 113 |
| 7 | 46 | 37 | (80.4) | 9 | (19.6) | 16 | (36.4) | 28 | (63.6) | 44 |

**Table S4**. Proportions of *PepNet* predictions on GRAMPA (N=1,707) and non-GRAMPA (N=1,180) validation sets and across folds.

| | GRAMPA | | | | | Non-GRAMPA | | | | |
|---|---|---|---|---|---|---|---|---|---|---|
| *Fold* | *Total* | *TP* | *(%)* | *FN* | *(%)* | *FP* | *(%)* | *TN* | *(%)* | *Total* |
| 1 | 1119 | 1009 | (90.2) | 110 | (9.6) | 119 | (28.0) | 306 | (72.0) | 425 |
| 5 | 261 | 234 | (89.7) | 27 | (10.3) | 72 | (18.6) | 315 | (81.4) | 387 |
| 6 | 244 | 199 | (81.6) | 45 | (18.4) | 50 | (16.6) | 252 | (83.4) | 302 |
| 7 | 83 | 73 | (88.0) | 10 | (12.0) | 20 | (30.3) | 46 | (69.7) | 66 |

**Table S5**. Proportions of *PepNet* predictions on GRAMPA (N=1,012) and non-GRAMPA (N=604) validation sets and across folds – leakage-free.

| | GRAMPA | | | | | Non-GRAMPA | | | | |
|---|---|---|---|---|---|---|---|---|---|---|
| *Fold* | *Total* | *TP* | *(%)* | *FN* | *(%)* | *FP* | *(%)* | *TN* | *(%)* | *Total* |
| 1 | 700 | 596 | (85.1) | 104 | (14.9) | 107 | (45.0) | 131 | (55.0) | 238 |
| 5 | 124 | 98 | (79.0) | 26 | (21.0) | 67 | (32.0) | 142 | (68.0) | 209 |
| 6 | 142 | 100 | (70.4) | 42 | (29.6) | 50 | (44.2) | 63 | (55.7) | 113 |
| 7 | 46 | 36 | (78.3) | 10 | (21.7) | 19 | (43.2) | 25 | (56.8) | 44 |
